# Adherent-to-suspension transition promotes melanoma metastatic dissemination

**DOI:** 10.1101/2024.11.26.625571

**Authors:** Dong Ki Lee, Jongwook Oh, Soyeon Lee, Hyunbin D. Huh, Yujin Sub, Hyun Woo Park, Heon Yung Gee

**Affiliations:** Department of Pharmacology, Yonsei University College of Medicine, Seoul, Republic of South Korea; Department of Pharmacology, Graduate School of Medical Science, Brain Korea 21 Project, Yonsei University College of Medicine, Seoul, 03722, Republic of Korea; Department of Biochemistry, College of Life Science and Biotechnology, Brain Korea 21 Project, Yonsei University, Seoul, 03722, Republic of Korea; Woo Choo Lee Institute for Precision Drug Development

**Keywords:** melanoma, adherent-to-suspension transition, circulating tumor cell, anchorage dependence

## Abstract

Melanoma is a highly metastatic skin cancer that often evades current therapeutic strategies primarily because of the complex mechanisms involved in metastasis. This study investigated the role of adherent-to-suspension transition (AST) in melanoma and its potential to facilitate metastasis by reprogramming cellular anchorage dependency. Using melanoma models, we demonstrated that the AST factors IKZF1, IRF8, and NFE2 are crucial for reprogramming, influencing gene expression related to cell adhesion and survival. Notably, our results highlight that AST factor expression undergoes dynamic changes during metastasis, which can be reversed in circulating tumor cells. Our findings revealed that AST contributes to the increased metastatic potential and invasiveness of melanoma cells and underscores its role independent of the epithelial-to-mesenchymal-like transition pathways. Based on the results, we highlight the potential of targeting AST mechanisms to develop new therapeutic strategies against metastasis.

## Introduction

Metastasis, the dissemination of cancer cells to organs distant from their origin, is the main barrier to effective cancer treatment and is the leading cause of cancer-related mortality (1,2). This process encompasses a series of biological events, including invasion of cancer cells at the primary site, entry into the bloodstream, survival within the circulatory system, exit from the bloodstream into adjacent tissues, and colonization and proliferation at a new site (3). While primary tumors can often be cured with local therapies, such as surgery and radiation, metastatic cancer is a systemic disease affecting multiple organs, culminating in the patient’s death (1,4).

Melanoma is a skin cancer with the highest mortality rate and presents formidable therapeutic challenges owing to its propensity for metastasis. When detected at an early stage, when the cancer is confined to its primary site, surgical resection offers high cure rates, with a 5-year survival rate exceeding 99% (5). However, the prognosis drastically changes once melanoma metastasizes to distant organs because its resistance to available therapies reduces the 5-year survival rate to approximately 30% (5). Hence, elucidating the mechanisms underlying melanoma metastasis is important for advancing melanoma treatment strategies.

Circulating tumor cells (CTCs) are essential precursors of metastasis (6). CTCs participate in the intermediate phases of the metastatic cascade. Recent technological advancements have enabled the detection and isolation of scarce CTCs (7,8). Studies have revealed the unique biological characteristics of CTCs, including the presence of CTC clusters and their interaction with the immune system (9,10). Nonetheless, how CTCs originate from the primary tumor and what prompts their detachment require further clarification. The epithelial-to-mesenchymal transition (EMT) involves a transformation whereby epithelial cells lose their characteristic adhesion and polarity, adopting traits that enable migration and invasion, contributing to the cancer’s ability to invade, metastasize, develop resistance to drugs, and maintain stemness and plasticity (11,12). Although melanoma is classified as a non-epithelial cancer, accumulating evidence indicates that the adoption of mesenchymal-like features enhances its tendency to metastasize and resistance to therapy (13,14). Since epithelial and mesenchymal states are typically associated with adherent cell types, the possibility of directly reprogramming the cell anchorage dependency from adherent to suspension states must be analyzed. A recent study demonstrated that a combination of specific hematopoietic transcriptional regulators can trigger the adherent-to-suspension transition (AST), changing anchorage dependency in solid tumors (15,16). In this study, we identified and characterized the functions of transcriptional regulators implicated in melanoma metastasis.

## Materials and Methods

### Cell culture

Cells were maintained in a humidified incubator at 5% CO_2_ and 37 °C. HEK293T, A375 iRFP, SK-MEL-28, IGR1, and B16F10 cells were cultured in Dulbecco’s modified Eagle’s medium (HyClone, SH30022.01) supplemented with 10% fetal bovine serum (FBS; HyClone, SV30207.02) and 1% penicillin/streptomycin (Invitrogen, 15140122). G361 cells were cultured in McCoy’s medium (Gibco, 16600082), supplemented with 10% fetal bovine serum (FBS) and 1% penicillin/streptomycin. No cell lines used in this study were found in the databases of commonly misidentified cell lines maintained by the International Cell Line Authentication Committee and NCBI BioSample database. The cell lines were tested and confirmed to be mycoplasma-free.

### Viral infection

HEK293T cells were transfected with plasmids encoding pMD2G and psPAX2, along with constructs cloned into a lentiviral vector, using Lipofectamine 2000 Transfection Reagent (11668019; Invitrogen), according to the manufacturer’s protocol. Media containing viral particles were harvested at 48 h post-transfection, passed through a 0.45-μm filter, supplemented with 8 mg/mL polybrene, and used for infection. Twenty-four hours after infection, transduced cells were incubated with fresh medium for another 24 h and selected using puromycin.

### Quantitative real-time PCR analysis

Cells were harvested for RNA extraction using a Hybrid-R™ kit (GeneAll^®^, 305-101). RNA samples were reverse-transcribed to complementary DNA using RNA-to-cDNA EcoDry Premix (639549; Takara). qRT-PCR was performed using a TB Green^®^ Premix Ex Taq™ kit (Takara, RR420B) and Quant Studio 3 (Applied Biosystems). The following primers were used for qPCR: IKZF1, 5′-TTTCAGGGAAGGAAAGCCCC-3′ (forward) and 5′-CTCCGCACATTCTTCCCCAT-3′ (reverse); IRF8, 5′-AGCATGTTCCGGATCCCTTG-3′ (forward) and 5′-CGGTCCGTCACTTCCTCAAA-3′ (reverse); NFE2, 5′-ACAGCTGTCCACTTCAGAGC-3′ (forward) and 5′-TGAGCAGGGGCAGTAAGTTG-3′ (reverse) for human cell lines.

### Orthotopic mouse model experiments

All animal experimental protocols were reviewed and approved by the Institutional Animal Care and Use Committee of Yonsei University. All mice were handled in accordance with the Guidelines for the Care and Use of Laboratory Animals. NOD/Shi-scid IL2 receptor gamma-null (NOG) mice were purchased from KOATECH, Inc., Republic of Korea. For the A375 cell footpad implantation model, 1[×[10^7^ iRFP-positive A375 melanoma cells were subcutaneously implanted into the footpads of 5-week-old male mice. NOG mice used in these experiments had identical genotypes and genetic backgrounds. C57BL/6 mice were purchased from Orient Bio, Inc. (South Korea). For the B16F10 cell footpad implantation model, 5[×[10^5^ GFP-positive B16F10 melanoma cells were subcutaneously implanted into the footpads of 7-week-old male mice. None of the experiments performed in this study surpassed the size limit of the tumors, that is, the volume did not exceed 2 cm^3^.

### Melanoma primary tumor and lung metastasis tissue dissociation

iRFP-positive A375 and GFP-positive B16F10 cells were isolated from primary tumor sites and lungs. Primary tumor and metastasized lung tissues were digested in 5 mL of enzyme buffer containing 0.2 mg/mL collagenase type-II (Worthington, LS004174), 0.1 mg/mL DNase I (Roche, 11284932001), and 0.8 mg/mL dispase (Gibco, 17105041) at 37 °C for 45 minutes. After complete digestion, the resulting cell suspension was filtered through a 40 μm cell strainer and washed. RBC lysis was then performed in suspension using ACK lysis buffer (Gibco, A1049201) for 5 min at room temperature.

### Generation of single-cell RNA sequencing data from orthotopic mouse models

For the A375-derived orthotopic mouse model, single-cell RNA sequencing (scRNA-seq) libraries were created using the Chromium Next GEM Single Cell 3p RNA library v3.1 (10X Genomics) with a target of 10,000 cells per library according to the manufacturer’s suggestions. This droplet-based system uses barcodes (one for each cell) and unique molecular identifiers (UMIs, one for each unique transcript) to obtain a unique 3′-mRNA gene expression profile from every captured cell. All samples were sequenced using Illumina HiSeq X Ten.

### scRNA-seq analysis of primary tumor cells, CTCs, and metastatic tumor cells

The Illumina outputs of A375-derived mouse samples and B16F10-derived mouse samples were mapped to the human reference genome (GRCh38) and mouse reference genome (GRCm38/mm10), respectively, using Cell Ranger 6.0.0 (10X Genomics), respectively. Gene-wise read counts for genes with a minimum of 500 reads were exported from Cell Ranger to the Matrix Market format and imported into R using Seurat’s Read10X function. Each 10X library was individually checked for quality, and the cells were filtered to ensure good gene coverage, a consistent range of read counts, and few mitochondrial reads. At least 500 detected genes were required for each cell, although the lower limit was reduced to 200 or 300 for some libraries. No more than 10% of mitochondrial reads were allowed per cell. Cells with exceptionally high read counts or detected genes were filtered to minimize the occurrence of doublets. After quality filtering, 19,419 cells from the A375-derived model (11,899 primary tumor cells, 2,838 CTCs, and 4,682 metastatic tumor cells) and 3,898 cells from the B16F10-derived model (1,129 primary tumor cells, 1,139 CTCs, and 1,630 metastatic tumor cells) were subjected to further analysis. Statistical analyses of the 10X data were conducted using the Seurat software package (v4.1.0) for R (17).

The samples were combined by merging them in Seurat. The cell clusters were identified using the default Louvain clustering algorithm implemented in Seurat. Default Seurat function settings were used, and 1:10 principal component dimensions were used for all the dimension reduction and integration steps. The cluster resolution was set to 1.0 unless otherwise stated. The RunUMAP random seed was set to 1,000 to ensure reproducibility.

### Immunocytochemistry of AST factors

For immunofluorescence staining, the primary tumors were fixed in 4% paraformaldehyde overnight, paraffin-embedded, and sectioned. Paraffin-embedded sections of primary tumors were rehydrated using a series of processes with xylene and different concentrations of alcohol and washed with phosphate-buffered saline (PBS). Samples were permeabilized with 0.15% Triton X-100 in PBS and blocked with 5% goat serum in PBS. The cells were then incubated at 4 °C overnight with the following primary antibodies at 1:100 dilutions: anti-Ikzf1 (rabbit, D6N9Y, Cell Signaling Technology), anti-CD31 (mouse, TLD-3A12, Thermo Fisher Scientific), and anti-GFP (chicken, ab13970, Abcam). After several washes, samples were incubated with secondary antibodies for 1 h at room temperature. The samples were then mounted with a fluorescent mounting medium, and immunofluorescent images were acquired using an LSM700 or LSM780 confocal microscope.

### Statistical analysis

All quantitative data were obtained from at least three independent biological replicates. Data are presented as mean [±[standard deviation unless otherwise noted in the figure legends. Statistical differences between two groups were examined using a two-tailed, unpaired Student’s *t*-test and one-way analysis of variance with Bonferroni correction for multiple comparisons. Statistical tests were performed using the GraphPad Prism 8.0 software (GraphPad Software, CA, USA). Two-sided *p*-values less than 0.05 were considered significant. No statistical methods were used to determine the sample size. The sample size was based on previous experience with experimental variability. Blinding was performed wherever possible during all sample analyses by coding sample identities during data collection, and analysis was performed by an observer without knowledge of or access to experimental conditions.

### Data availability

Single-cell RNA-seq data were deposited in the Korean BioData Station (K-BDS, https://kbds.re.kr) with accession number KAP240742. Spatial transcriptomic data of human melanoma tissue were generated using the Spatial Gene Expression Dataset by Space Ranger v2.0.0, 10X Genomics (2022, July 13). Further inquiries can be directed to the corresponding author.

## Results

### Induction of AST factors in disseminated melanoma cancer CTCs

To identify the key factors required for reprogramming anchorage dependency, we investigated the *in vivo* significance of the AST paradigm in the metastatic spread and survival of CTCs using mouse models. We injected iRFP-expressing A375 melanoma cells harboring the BRAF V600E mutation into the footpads of mice. After confirming metastasis, we performed a comparative transcriptomic analysis across three distinct stages of hematogenous melanoma dissemination, using iRFP-labeled A375 melanoma cells isolated from the primary tumor, whole blood, and metastatic tumor (**Fig. 1A**). To ensure the purity of the melanoma cells, we analyzed scRNA-seq data from CD45-negative cells and found that the majority of cells were positive for melanocyte markers (Supplementary Fig. S1A and S1B). After filtering the cells using quality control measures and correcting for batch effects, we visualized the sequencing data of 11,899 primary tumor cells, 2,838 CTCs, and 4,682 metastatic tumor cells using Uniform Manifold Approximation and Projection (UMAP) (**Fig. 1B**; Supplementary Fig. S1C). Given the unique ability of CTCs to spread and persist within the bloodstream, we anticipated that CTCs would possess transcriptional signatures distinct from those of primary or metastatic tumor cells. Despite partial overlap, clusters of CTCs separated from the primary or metastatic tumor cells were evident (**Fig. 1B**), suggesting transcriptional reprogramming during dissemination.

**Fig. 1.**
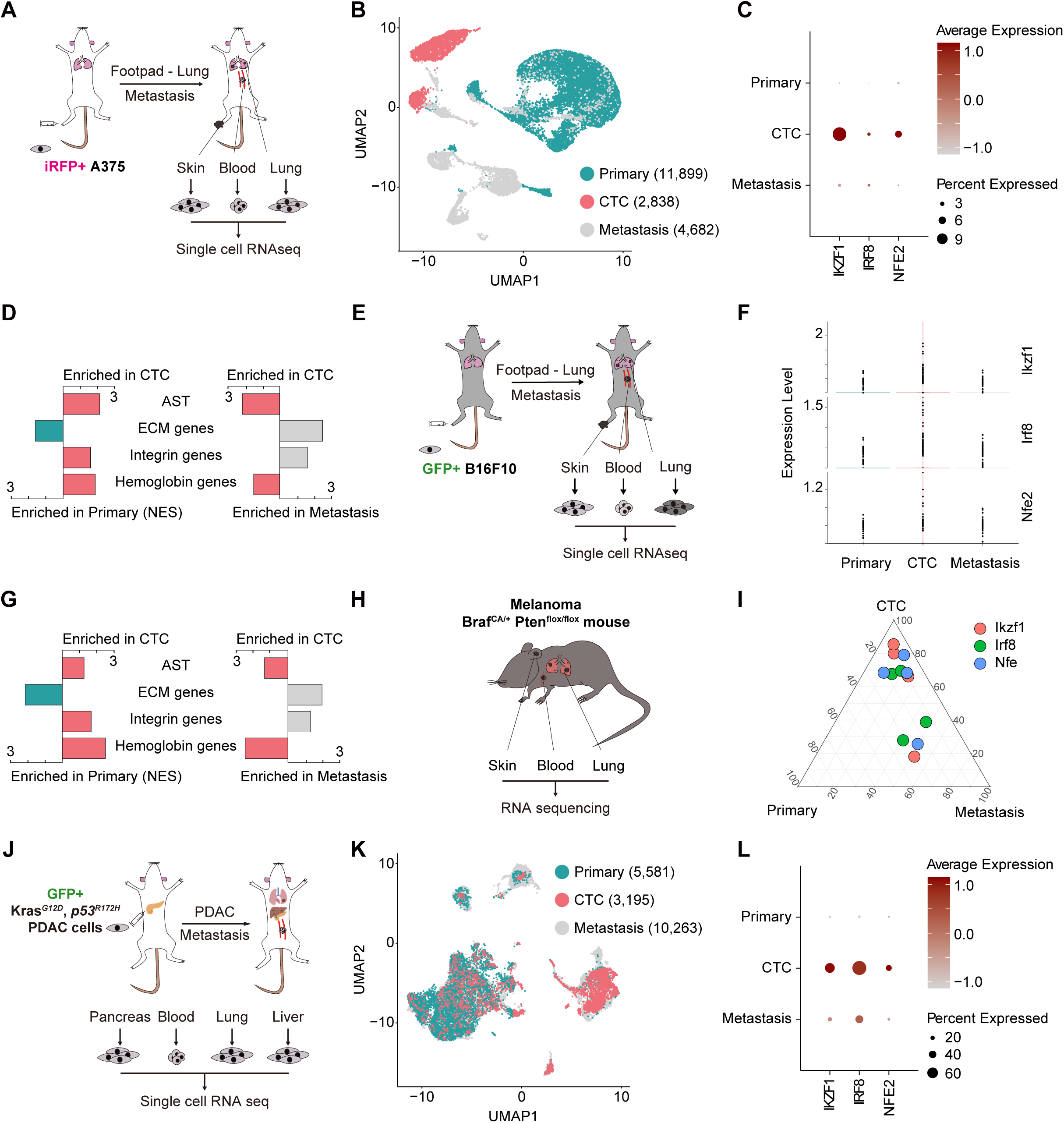
Induction of AST factors in disseminated CTCs of various *in vivo* models. **A**, Schematic overview of the isolation and scRNA-seq analysis of A375-derived primary tumor cells, CTCs, and metastatic tumor cells from an orthotopic melanoma metastasis model. iRFP-positive cells were injected into the footpads of NOG mice, and tumor cells were subsequently isolated using FACS. **B**, UMAP embedding of the analyzed transcriptomes of 11,899 primary tumor cells (blue), 2,838 CTCs (red), and 4,682 metastatic tumor cells (gray). **C**, Dot plot showing the average expression levels of IKZF1, IRF8, and NFE2 in primary tumor cells, CTCs, and lung metastatic tumor cells, respectively. **D**, Bar plot displaying the GSEA results for AST, ECM, integrin, and hemoglobin genes among A375-derived primary tumor cells, CTCs, and lung metastasized tumor cells. **E**, Schematic overview of the isolation and scRNA-seq analysis of B16F10-derived primary tumor cells, CTCs, and metastatic tumor cells from an orthotopic melanoma metastasis model. GFP-positive cells were injected into the footpads of C57BL/6N mice, and tumor cells were subsequently isolated using FACS. **F**, Violin plot showing the average expression levels of three factors, Ikzf1, Irf8, and Nfe2, in primary tumor cells, CTCs, and lung-metastasized tumor cells. **G**, Bar plot displaying the GSEA results for AST, ECM, integrin, and hemoglobin genes in B16F10-derived primary tumor cells, CTCs, and lung-metastasized tumor cells. **H**, Schematic overview of the inducible B-RAF–PTEN-driven melanoma mouse model. **I**, Ternary plot illustrating AST gene expression in paired primary tumors, CTCs, and lung metastatic tumors. **J**, Schematic overview of the isolation and scRNA-seq analysis of KRAS, TP53 mutant pancreatic cancer cell line-derived primary tumor cells, CTCs, and metastatic tumor cells from an orthotopic pancreatic cancer metastasis model. **K**, UMAP embedding of analyzed transcriptomes of 5,581 primary tumor cells (blue), 3,195 CTCs (red), and 10,263 metastatic tumor cells (gray). **L**, Dot plot showing the average expression levels of IKZF1, IRF8, and NFE2 in primary tumor cells, CTCs, and metastasized tumor cells.

Previous research has identified IKZF1, NFE2, BTG2, and IRF8 as key factors that promote anchorage-independent growth in solid tumors (15). The AST mechanism has been delineated into two main stages: the first involves spontaneous cell-matrix dissociation due to YAP/TEAD pathway suppression, which leads to decreased expression of extracellular matrix (ECM)-integrin genes, and (second encompasses the development of resistance to anoikis following the activation of hemoglobin genes (15). Consistent with these findings, the levels of IKZF1, IRF8, and NFE2 expression were higher in CTCs than in primary tumor cells and then reverted back to their initial levels in metastatic tumor cells (**Fig. 1C**). Further investigation revealed distinctive transcriptional behaviors in tumor cells at primary, CTCs, and metastatic sites. Specifically, the generation of CTCs showed a reduction in the expression of ECM genes and an increase in the expression of hemoglobin genes, consistent with earlier observations (**Fig. 1D**) (15). This pattern was reversed during the formation of metastatic sites (**Fig. 1D**).

In our previous study (15), we examined the AST phenomenon in a syngeneic mouse model of melanoma using GFP-labeled B16F10 cells (**Fig. 1E**) that lacked the BRAF mutation. Analysis of scRNA-seq data of 1,129 primary tumor cells, 1,139 CTCs, and 1,630 metastatic tumor cells (Korea BioData Station accession number KAP230547; ref. (15)), we observed a consistent pattern of AST gene expression across the three groups, demonstrating the plasticity of AST (**Fig. 1F**). These findings in mouse models using two different melanoma cell lines suggest that the AST phenomenon is independent of the BRAF V600E mutation. Consistent dynamics were observed in the expression of ECM and hemoglobin genes throughout the metastatic cascade (**Fig. 1G**). These distinctive transcriptional behaviors of CTCs were replicated in a genetically engineered mouse model of melanoma (**Fig. 1H**) (18). When examining paired samples, there was a noticeable increase in AST factor expression coinciding with the generation of CTCs, which then decreased during the formation of metastases (**Fig. 1I**; Supplementary Fig. S3A). A similar increase in AST factor expression during CTC formation was noted in an orthotopic mouse model of pancreatic cancer (**Fig. 1J**) (19). Within the published dataset, including 5,581 primary tumor cells, 3,195 CTCs, and 10,263 metastatic tumor cells (**Fig. 1K**), there was a pronounced upregulation of AST factors in CTCs relative to cells from primary or metastatic tumors (**Fig. 1L**). Constant trends in the expression of ECM and hemoglobin genes were identified along the metastatic cascade in both the genetically engineered mouse model of melanoma and orthotopic mouse model of pancreatic cancer (Supplementary Fig. S3B and S3C). This finding suggests that the influence of AST extends beyond that of melanoma.

### An AST continuum independent of EMT-like transition

Next, we aimed to elucidate the transcriptional relationship between AST and EMT-like transitions during melanoma dissemination. In CTCs, three AST factors showed increased expression, whereas EMT-like transition-linked genes, including AXL and ZEB1 (20), displayed inconsistent expression (**Fig. 2A** and **B**). Conversely, indicators of MET-like transitions such as MITF and ZEB2 (20) showed increased expression in CTCs (**Fig. 2A** and **B**). This suggests a distinction between AST- and EMT-like transition mechanisms in the progression of melanoma metastasis.

**Fig. 2.**
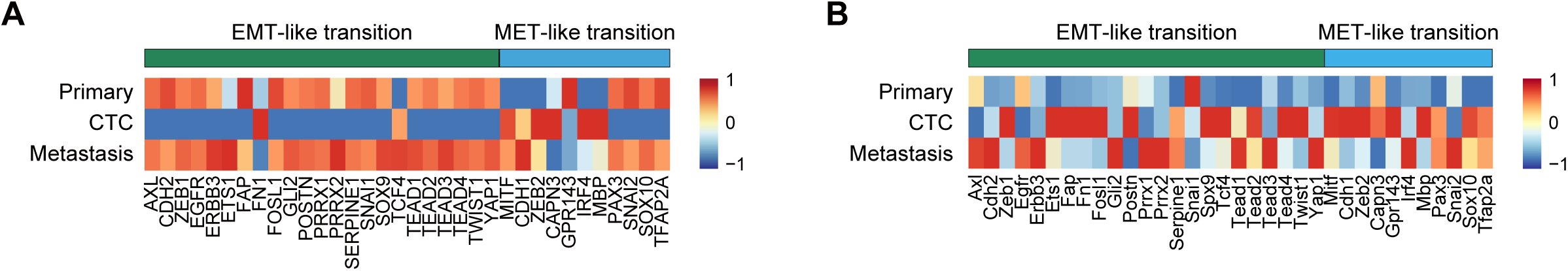
EMT-like transition during the metastatic cascade and a transcriptional AST continuum *in vivo*. **A**, Heatmap displaying the expression of EMT-like and MET-like transition genes in A375-derived primary tumor cells, CTCs, and lung-metastasized tumor cells. **B**, Heatmap displaying the expression of EMT-like and MET-like transition genes in B16F10-derived primary tumor cells, CTCs, and lung-metastasized tumor cells.

### AST expression contributes to clone aggressiveness

To determine whether the induction of AST factors increases the propensity for metastatic dissemination of melanoma cells, we conducted clonal phylogenetic analysis using our scRNA-seq data informed by copy number variation profiles (**Fig. 3A**) (21). Our findings indicated that cells within clone 1 harbored a higher proportion of metastatic tumor cells than primary tumor cells (**Fig. 3B**). Moreover, CTCs within clone 1 exhibited a pronounced increase in AST factor expression compared to CTCs from different clones (**Fig. 3C**). Therefore, we examined the transcriptional signatures of the clones. Notably, CTCs from clone 1 displayed a unique transcriptional profile, characterized by the reduced expression of ECM genes and genes associated with hypoxia and reactive oxygen species (ROS) pathways (**Fig. 3D**). Based on our observations, we postulated that CTCs generated via the induction of AST factors have an enhanced likelihood of metastasis.

**Fig. 3.**
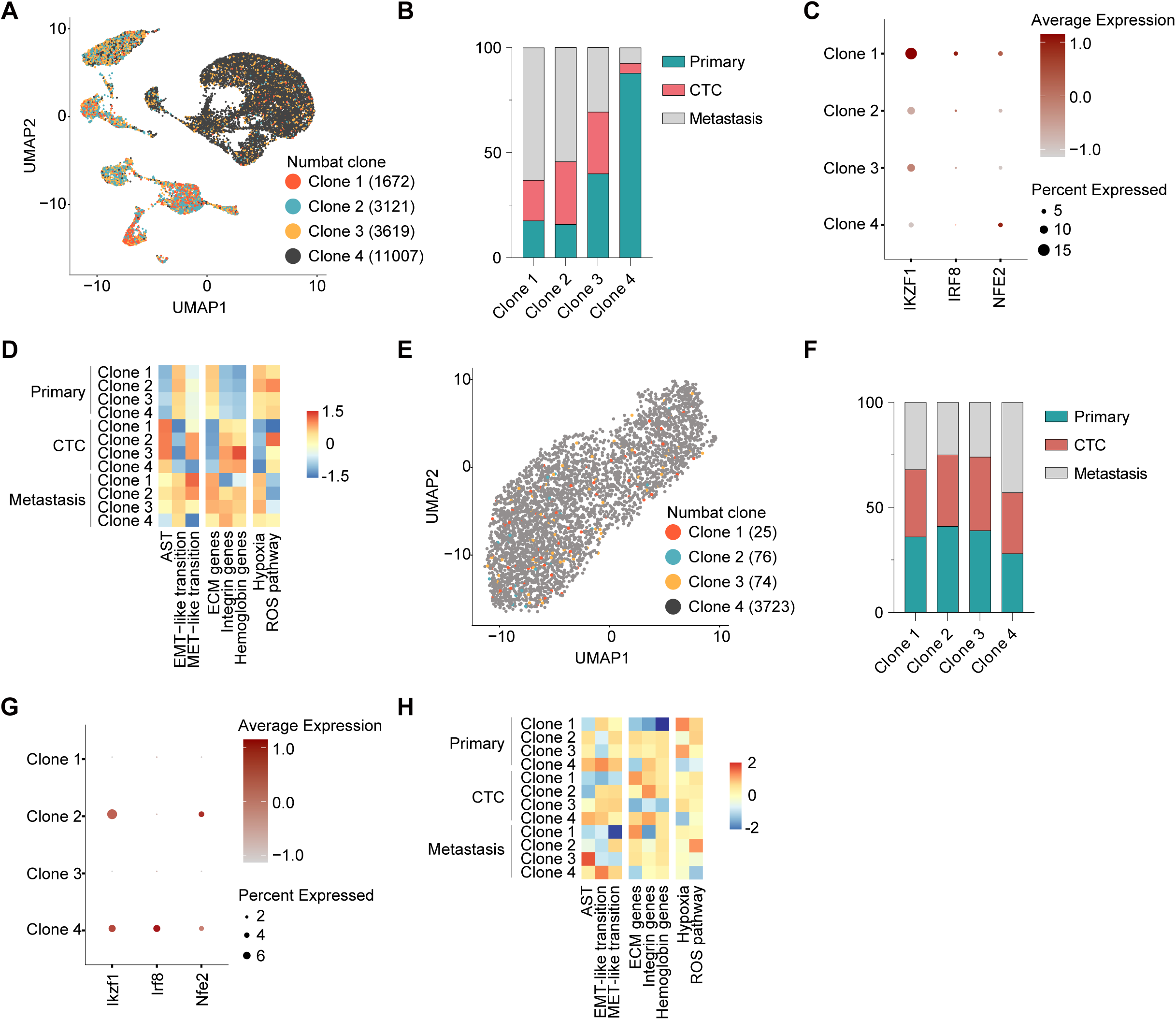
Impact of AST expression on the metastatic potential of tumor clones. **A**, UMAP embedding of transcriptomes from clonal phylogenetic analysis based on copy number variation profiles of A375-derived scRNA-seq data, showing 1,672 Clone 1 cells (red), 3,121 Clone 2 cells (blue), 3,619 Clone 3 cells (yellow), and 11,007 Clone 4 cells (black). **B**, Proportion of primary tumor cells (blue), CTCs (red), and metastatic tumor cells (gray) in each clone. **C**, Dot plot depicting the average expression levels of IKZF1, IRF8, and NFE2 in CTCs of each clone. **D**, Single-sample gene set enrichment analysis results for each clone classified according to the origin of the tumor cells. **E**, UMAP embedding of transcriptomes from clonal phylogenetic analysis based on copy number variation profiles of B16F10-derived scRNA-seq data, showing 25 clone 1 cells (red), 76 clone 2 cells (blue), 74 clone 3 cells (yellow), and 3,723 Clone 4 cells (gray). **F**, Proportion of primary tumor cells (blue), CTCs (red), and metastatic tumor cells (grey) in each clone. **G**, Dot plot showing the average expression levels of IKZF1, IRF8, and NFE2 in CTCs of each clone. **H**, Single-sample gene set enrichment analysis results for each clone classified by the origin of the tumor cells.

Subsequent validation using the B16F10 mouse model confirmed this hypothesis (**Fig. 3E**). Among the clones, clone 4 had the highest rate of metastasis compared with its primary tumor cells, with CTCs from this clone expressing the highest levels of AST (**Fig. 3F** and **G**). Additionally, CTCs from clone 4 also showed a decrease in the expression of genes related to hypoxia and ROS pathways, suggesting that these changes could be attributed to AST factors that promote the formation of CTCs in solid tumors (**Fig. 3H**).

### AST factors reprogram the anchorage dependency of adherent cells

Although we examined the effects of AST on altering anchorage dependency and the spread of metastases *in vivo*, we also investigated the expression of AST factors in adherent cell lines to further elucidate the underlying mechanisms. *In vitro* studies of metastasis have primarily focused on the ability of cells to migrate and invade distant organs. However, investigating how cancer cells are influenced during dissemination is crucial. Thus, we developed a new *in vitro* assay termed the “dissemination assay,” which effectively mimics the complete metastatic cascade, including migration from the original tumor into the bloodstream and metastasis in new organs (**Fig. 4A**). Given that high cell density often leads to the competitive displacement of latent metastatic cells (22), we hypothesized that a portion of cancer cells might detach from culture plates when subjected to prolonged high-cell-density environments. To explore this, we used various melanoma cell lines derived from humans and mice, including A375 and B16F10 cells, which have been used in *in vivo* models. We observed that a subset of cells spontaneously rounded up and detached from the plates (**Fig. 4B** and **C**). When these cells were transferred to a new plate, they reverted to their adherent state and reattached. Furthermore, qRT-PCR analysis of AST factors showed significantly higher expression in spontaneously suspended cells than in adherent cells (**Fig. 4D**).

**Fig. 4.**
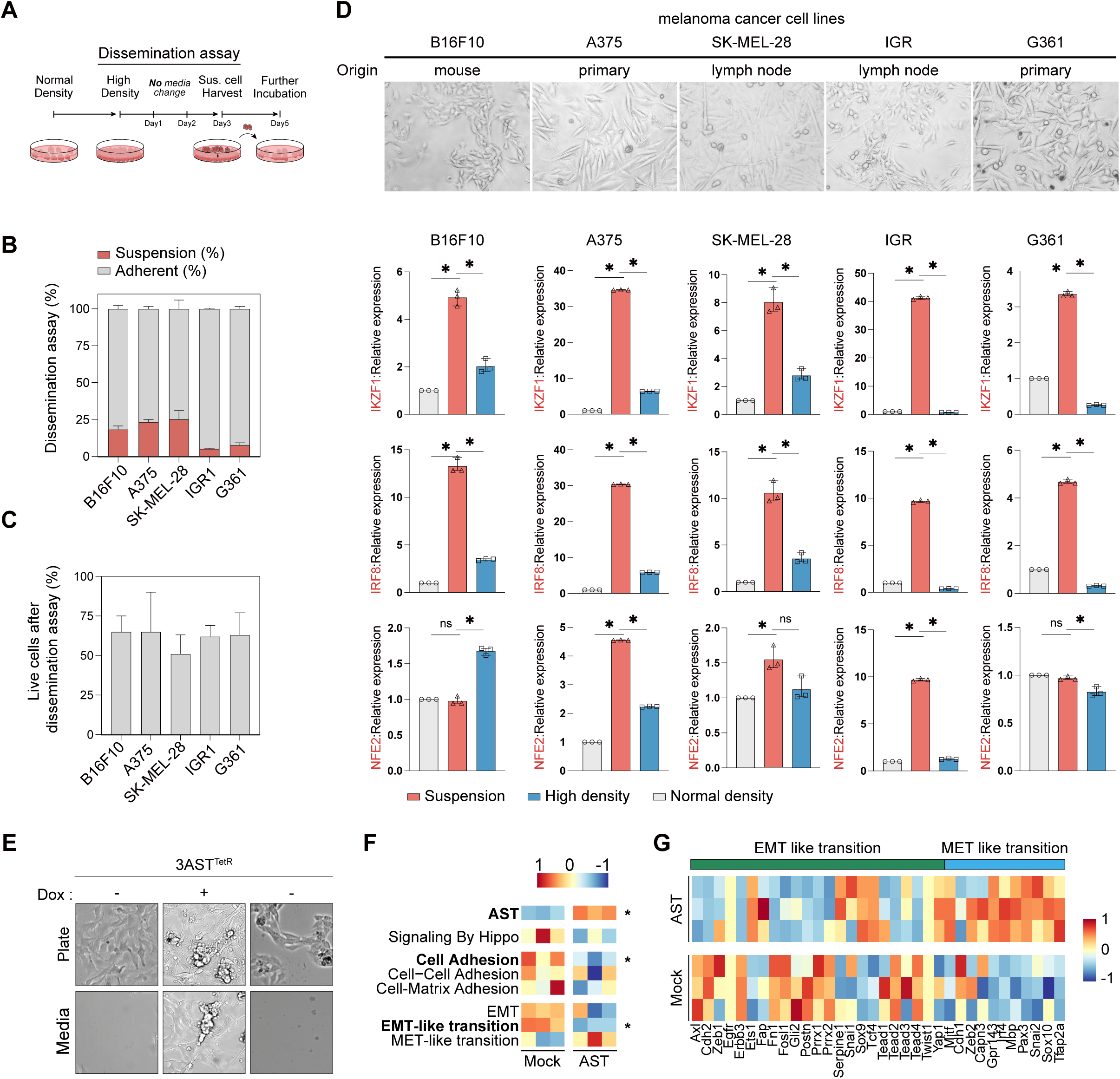
*In vitro* reproduction of AST factor induction during the metastatic cascade. **A**, Schematic overview of *in vitro* dissemination assay. Cells were seeded at a low density and incubated for 72–96 h until the cell density reached 100%. The cells were incubated for an additional 72 h, and spontaneously detached cells were harvested and subjected to further analyses. **B**, Relative efficiency of the dissemination assay, shown as the proportion of quantified suspension-state cells relative to quantified adherent-state cells. **C**, Proportion of live cells among the harvested suspension-state cells from various melanoma cell lines subjected to the dissemination assay. **D**, qRT-PCR analysis of AST expression at each phase of the dissemination assay in different melanoma cell lines. **E**, Consecutive reprogramming of anchorage dependency in B16F10-3AST^TetR^ cells by the addition and removal of doxycycline. Scale bars, 50 μm. **F**, Single-sample gene set enrichment analysis results for mock and AST-induced suspended cells. **G**, Heatmap of EMT-like and MET-like transition genes in mock- and AST-induced suspended cells.

To better understand the role of AST in regulating anchorage dependence, we engineered B16F10-3AST^TetR^ cells to undergo repeated cycles of doxycycline treatment (**Fig. 4E**). We observed that doxycycline administration caused these cells to detach, a process that was reversed by doxycycline withdrawal (**Fig. 4E**). Further exploration using RNA sequencing revealed upregulation of AST genes in AST-induced suspended cells (**Fig. 4F**). Additionally, these cells displayed reduced activity in the Hippo signaling pathway, consistent with previous results (**Fig. 4F**) (15). Importantly, we noted a reduction in cell adhesion, primarily attributable to decreased cell-matrix interactions rather than alterations in cell-cell adhesion (**Fig. 4F**).

AST-treated cells showed depletion of the Hallmark EMT pathway. Given that EMT-like transitions in melanoma do not correspond to conventional EMT, we compared the expression of EMT-like and MET-like transition genes in mock- and AST-induced suspended cells (**Fig. 4F** and **G**). EMT-like transition genes were predominantly upregulated in the control cells, whereas MET-like transition genes were highly expressed in AST-induced cells (**Fig. 4G**). Nonetheless, there was a discrepancy between the genes, resulting in non-significant gene set enrichment analysis (Supplementary Fig. S5D). Thus, our observations indicate that AST discretely reprograms the anchorage dependence from EMT-like or MET-like transition pathways.

## Discussion

Currently, the prevailing view of metastasis focuses heavily on genetic alterations. However, the recent discovery of common mutations between primary and metastatic tumors points to the role of additional nongenetic elements (23-26). This shift has sparked interest in cellular plasticity and the impact of environmental factors on cell traits, such as morphology and anchorage dependence during metastasis. This newly identified AST phenomenon clarifies a fundamental regulatory mechanism in CTC generation and cancer spread (15). Four key factors, IKZF1, NFE2, BTG2, and IRF8, drive this transition by altering the expression of genes associated with adhesion and survival. However, the application of AST in various types of cancers remains unclear. In the present study, we observed the induction of AST factors in CTCs disseminated from melanoma and pancreatic cancer models. We discovered that AST expression could be dynamically altered during CTC generation and the subsequent colonization of the metastatic cascade. Specifically, the downregulation of ECM genes and upregulation of hemoglobin genes, notably HBA1 and HBA2, were reversed during colonization at metastatic sites. Insights into AST plasticity may pave the way for new therapeutic approaches targeting the metastatic spread of cancer.

Although melanoma is not of epithelial origin, it undergoes plasticity similar to that observed in epithelial tumors (20). This EMT-like behavior, often termed phenotype switching, involves the dynamic reprogramming of melanoma cells, leading to a loss of melanocytic markers, increased invasiveness, metastasis, and treatment resistance. This process is regulated by key transcription factors such as MITF, ZEB1, and SOX10, and is more complex than that in epithelial cancers, with melanoma cells exhibiting a spectrum of intermediate phenotypes rather than a binary switch (13,14,20). However, the role of EMT-like transition in the process of CTC generation is poorly understood. This study introduces the concept of pseudo-AST, which reflects the spectrum of cells transitioning from an adherent to a suspended state within the primary tumor. Regarding pseudo-AST, the dynamics of the EMT-like and MET-like transition genes showed no clear patterns. This disorganization was also observed when pseudo-AST was used to analyze the primary tumor and CTC assemblies in the scRNA-seq data. Therefore, our findings suggest that AST reshapes anchorage dependence, which is distinct from EMT-like or MET-like transition pathways.

In our study, clones expressing AST exhibited the highest metastatic potential and aggressiveness among the studied tumor clones. This finding underscores the significant role of AST expression in enhancing the invasive capabilities of melanoma cells, thereby facilitating their spread and colonization of distant sites. The pronounced metastatic behavior of AST factor-positive clones can be attributed to their increased AST expression levels, which likely confer a competitive advantage in morphological transition after the loss of cell-matrix interaction and anoikis resistance within the bloodstream. Intriguingly, among the CTC clones, the AST factor-expressing clone exhibited comparatively lower enrichment of hypoxia and ROS pathways. This finding highlights the pivotal contribution of AST expression to the metastatic progression and inherent aggressiveness of melanoma and emphasizes the potential of targeting AST-related pathways as a therapeutic strategy to curb melanoma dissemination.

In conclusion, this study elucidated a complex mechanism of metastasis involving the AST phenomenon in melanoma, which is independent of the EMT-like and MET-like transition pathways. This study highlights the dynamic nature of AST expression during the metastatic cascades. Ultimately, insights into AST plasticity may lead to novel therapeutic strategies aimed at curbing the metastatic spread of cancer, emphasizing the importance of further exploration of this pathway.

## Acknowledgements

We thank all members of the Gee and Park laboratories for their comments on the manuscript and for their discussions.

