## Supplemental Figures 1-5 for "Adherent-to-suspension transition promotes melanoma metastatic dissemination"

### Contents

**1. Supplementary Figure 1.** scRNA-seq analysis of A375-derived orthotopic melanoma metastasis model.

**2. Supplementary Figure 2.** Single cell RNA-seq analysis of B16F10-derived orthotopic melanoma metastasis model.

**3. Supplementary Figure 3.** Gene expression profiles in various cancer metastasis mouse models.

**4. Supplementary Figure 4.** Clonal phylogenetic analysis based on copy number variation profiles of A375-derived scRNA-seq data highlights differences in the metastatic potential among tumor clones.

**5. Supplementary Figure 5.** Reprogramming of anchorage dependency in B16F10-3ASTTetR cells.

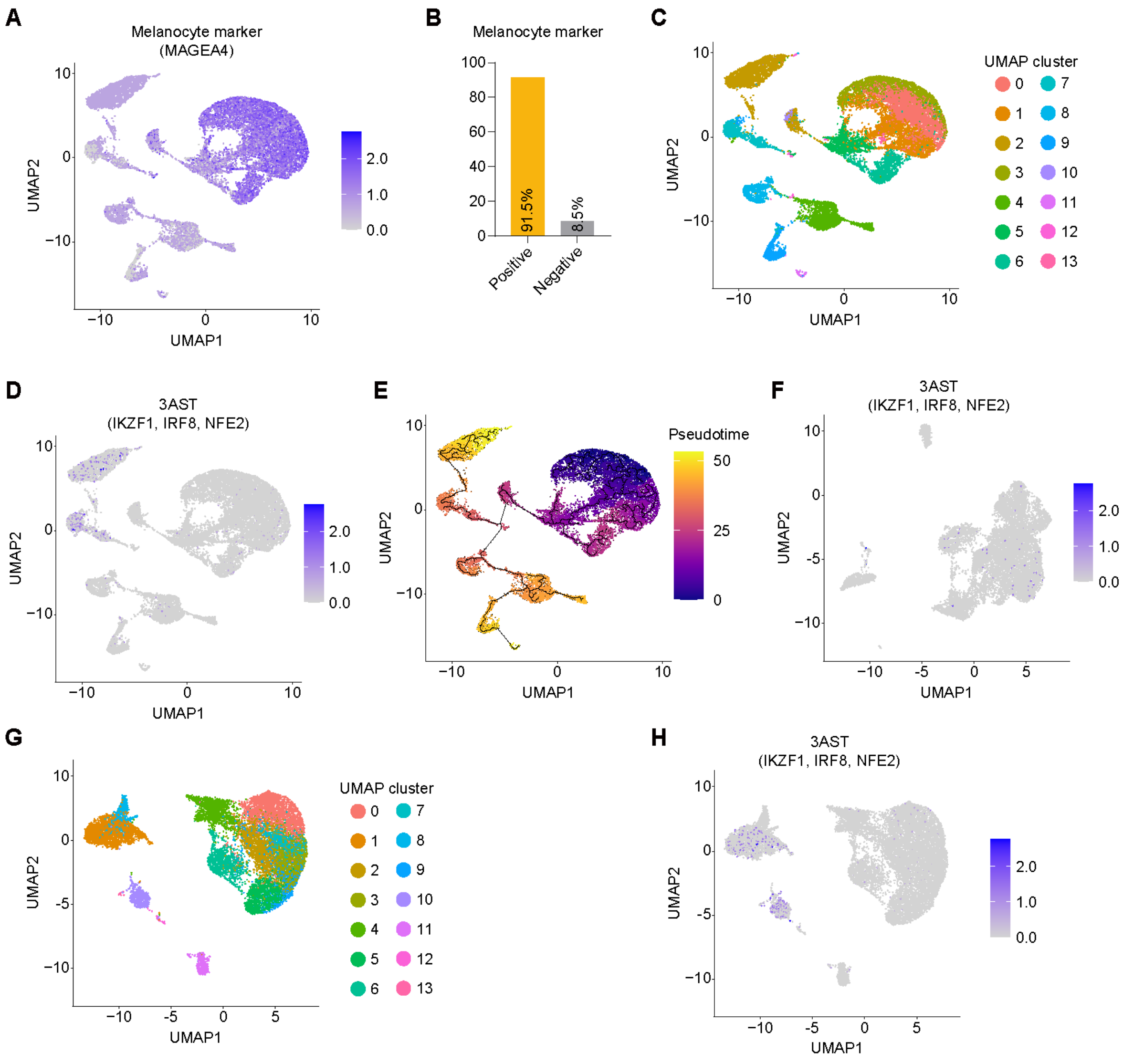


**Supplementary Figure 1. scRNA-seq analysis of A375-derived orthotopic melanoma metastasis model.**

**A**. Feature plot illustrating the expression levels of the melanocyte marker gene MAGEA4 across A375-derived primary tumor cells, CTCs, and metastatic tumor cells.

**B**. Bar plot depicting the proportion of cells that are positive or negative for the expression of the melanocyte marker gene.

**C**. UMAP visualization showing the clustering of cell populations identified within the A375-derived orthotopic melanoma metastasis model .

**D**. Feature plot illustrating the expression of AST genes (IKZF1, IRF8, and NFE2) across cell populations.

**E**. Monocle pseudo-time trajectory analysis of A375-derived tumor cells based on UMAP embeddings, showing progression of cell states.

**F**. Feature plot illustrating the expression of AST genes (IKZF1, IRF8, and NFE2) across A375-derived primary tumor cells.

**G**. UMAP visualization showing the clustering of cell populations identified within A375-derived primary tumor cells and CTCs.

**H**. Feature plot illustrating the expression of AST genes (IKZF1, IRF8, and NFE2) across A375-derived primary tumor cells and CTCs.

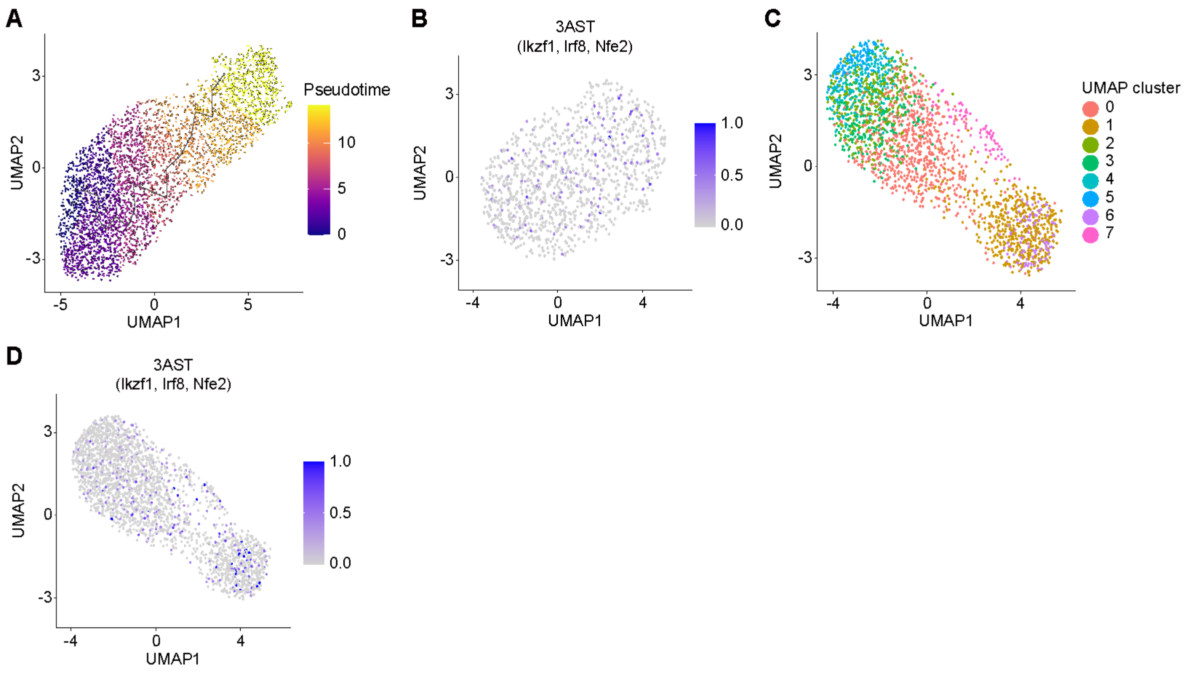


**Supplementary Figure 2. scRNA-seq analysis of B16F10-derived orthotopic melanoma metastasis model.**

**A**. Monocle pseudo-time trajectory analysis of B16F10-derived tumor cells based on UMAP embeddings, showing progression of cell states.

**B**. Feature plot illustrating the expression of AST genes (Ikzf1, Irf8, and Nfe2) across B16F10-dervied primary tumor cells.

**C**. UMAP visualization showing the clustering of cell populations identified within B16F10-derived primary tumor cells and CTCs..

**D**. Feature plot illustrating the expression of AST genes (Ikzf1, Irf8, and Nfe2) across B16F10-dervied primary tumor cells and CTCs.

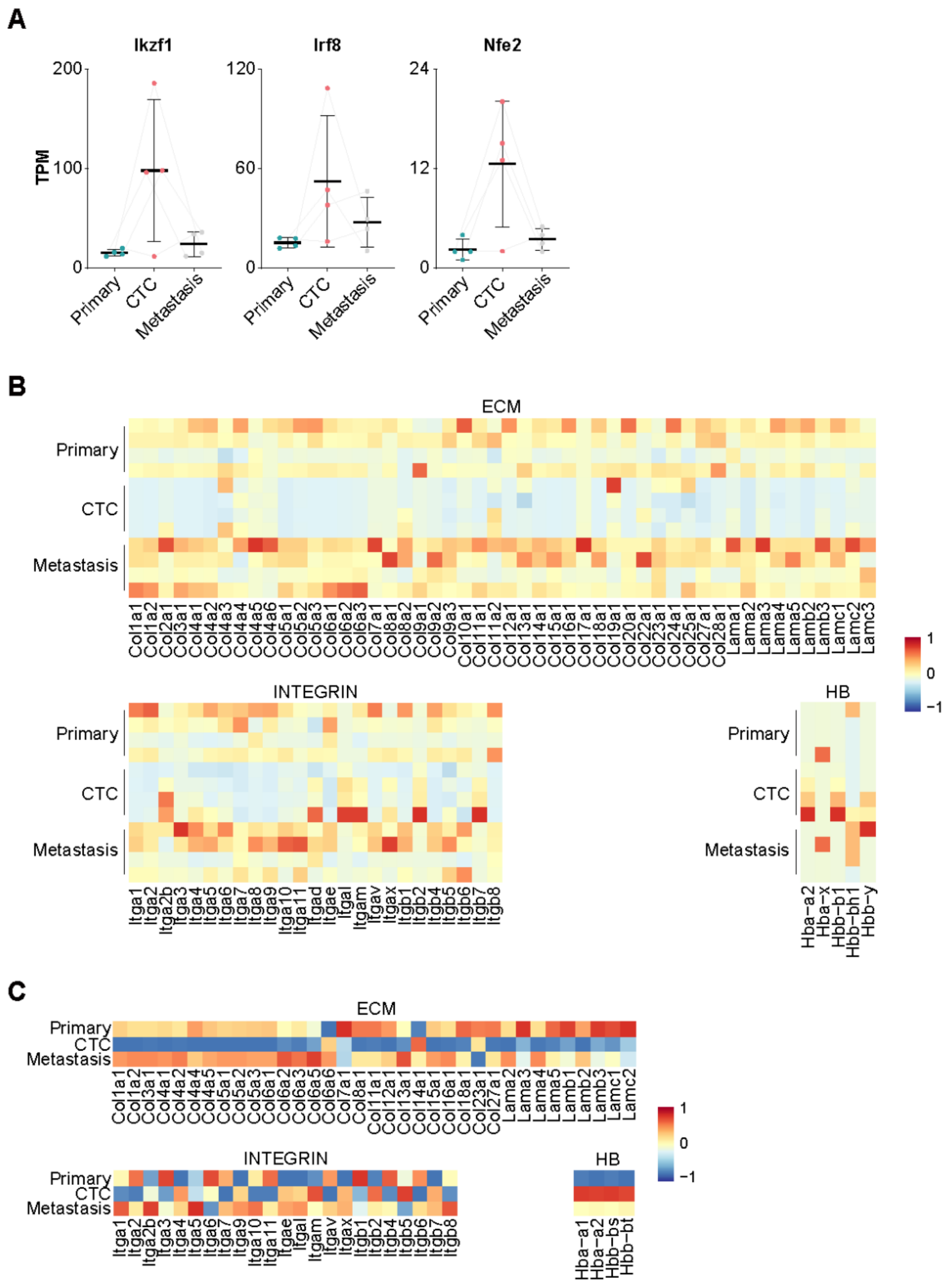


**Supplementary Figure 3. Gene expression profiles in various cancer metastasis mouse models.**

**A**. Dynamics of AST gene expression (Ikzf1, Irf8, and Nfe2) across matched primary tumors, CTCs, and metastatic tumor samples from the inducible BRAF–PTEN-driven melanoma mouse model.

**B**. Heatmap displaying the expression of ECM, Integrin, and Hemoglobin genes across primary tumors,

CTCs, and metastatic tumor samples in the inducible BRAF–PTEN-driven melanoma mouse model.

**C**. Heatmap displaying the expression of ECM, Integrin, and Hemoglobin genes across primary tumors, CTCs, and metastatic tumors based on pseudobulk differential expression analysis in the KRAS, TP53-mutant pancreatic cancer cell line-derived orthotopic pancreatic cancer metastasis model.

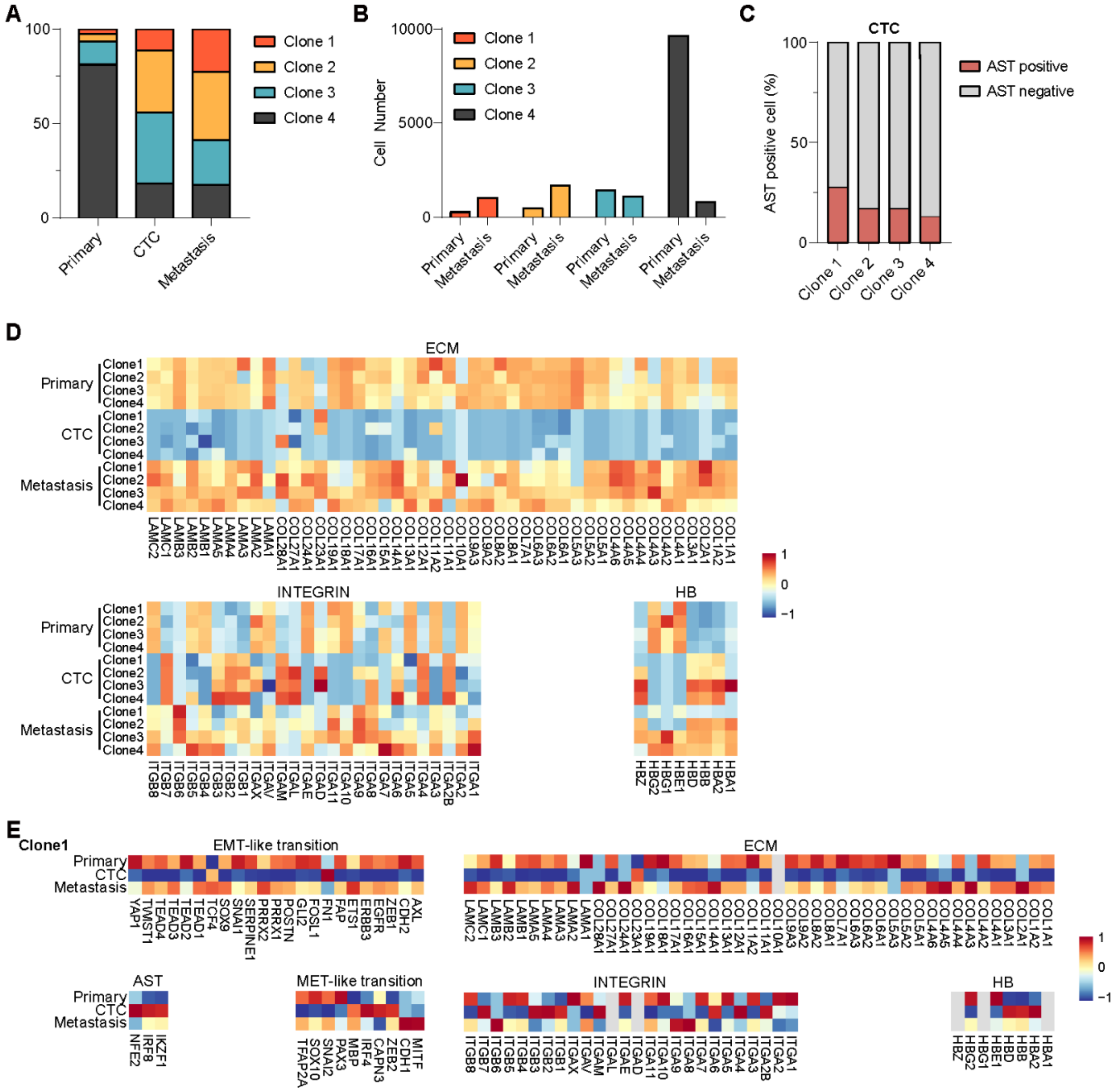


**Supplementary Figure 4. Clonal phylogenetic analysis based on copy number variation profiles of A375-derived scRNA-seq data highlights differences in the metastatic potential among tumor clones.**

**A**. Stacked bar plot showing the distribution of clones (Clone 1 to Clone 4) across A375-derived primary tumor cells, CTCs, and metastatic tumor cells.

**B**. Bar plot indicating the cell count of each clone in primary tumor cells, CTCs, and metastatic tumor cells, highlighting clonal dynamics across conditions.

**C**. Bar plot indicating the proportion of AST-positive and AST-negative cells within each clone in CTCs.

**D**. Heatmap displaying the expression of ECM, Integrin, and Hemoglobin genes across clones in primary tumors, CTCs, and metastatic samples.

**E**. Clone-specific heatmap illustrating the expression patterns of EMT-like transition, MET-like transition, AST, ECM, Integrin, and Hemoglobin genes in Clone 1 across primary tumors, CTCs, and metastatic samples, showing gene expression changes associated with metastatic progression.

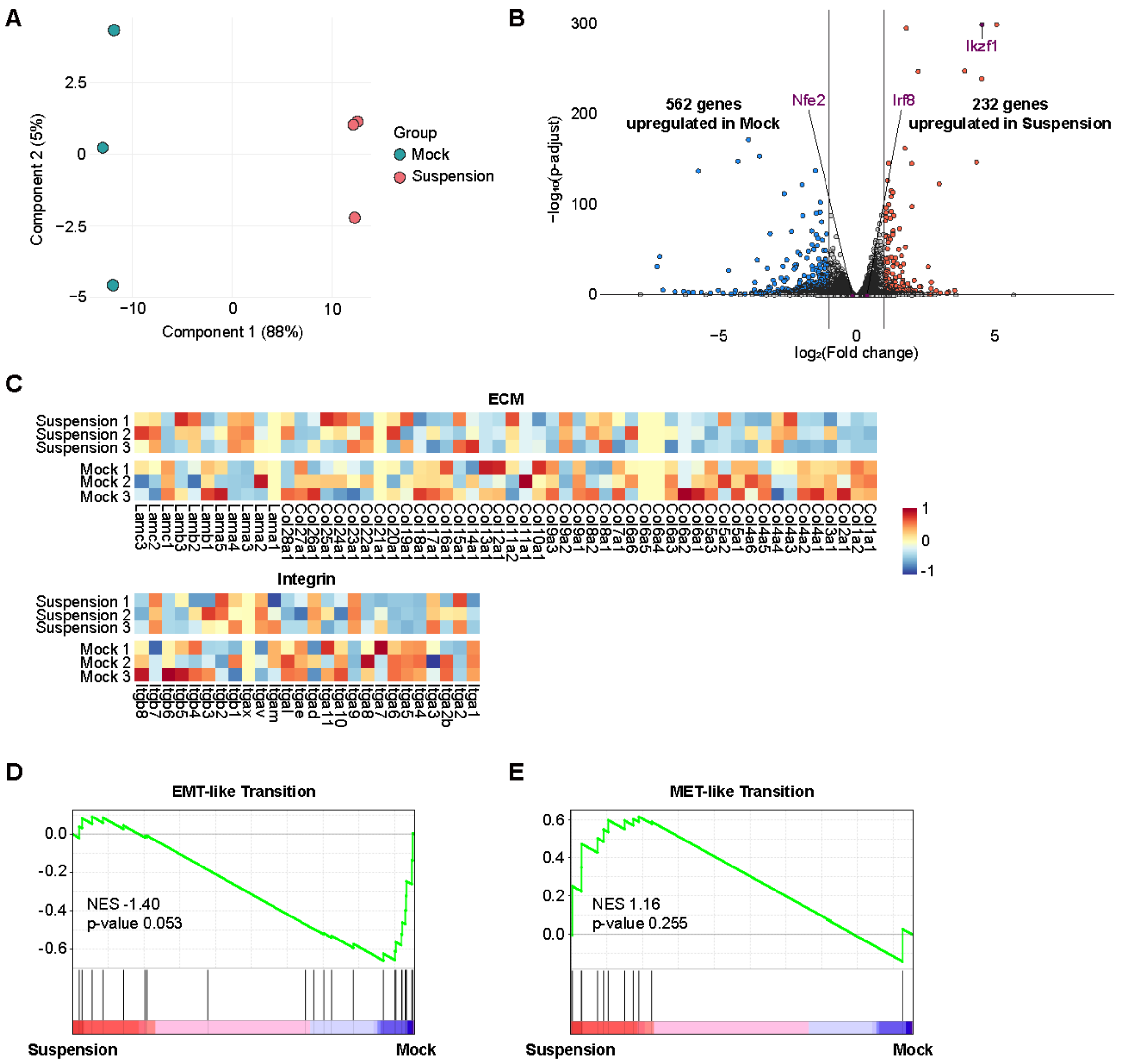


**Supplementary Figure 5. Reprogramming of anchorage dependency in B16F10-3AST$^{TetR}$ cells.**

**A**. Principal component analysis plot illustrating the clustering of suspension and mock samples, highlighting the distinction between the two groups.

**B**. Volcano plot depicting differentially expressed genes between suspension and mock groups, with Ikzf1 among the three AST genes showing significant upregulation in suspension conditions.

**C**. Heatmap displaying the expression of ECM, Integrin, and Hemoglobin genes across suspension and mock conditions.

**D-E**. Gene set enrichment analysis plots for EMT-like (D) and MET-like (E) transition gene sets comparing suspension and mock groups. The analysis indicates a trend toward EMT-like transition enrichment in the mock group and MET-like transition enrichment in the suspension group, though the results are not statistically significant due to variability among genes.
